# Using global t-SNE to preserve inter-cluster data structure

**DOI:** 10.1101/331611

**Authors:** Yuansheng Zhou, Tatyana O Sharpee

## Abstract

The t-distributed Stochastic Neighbor Embedding (t-SNE) method is one of the leading techniques for data visualization and clustering. This method finds lower dimensional embeddings of data points while minimizing distortions in distances between neighboring data points. By construction, t-SNE discards information about large scale structure of the data. We show that adding a global cost function to the t-SNE cost function makes it possible to cluster the data while preserving global inter-cluster data structure. We test the new “global t-SNE” (g-SNE) method on one synthetic and two real data sets on flowers and human brain cells which have significant and meaningful global structures. In all cases, g-SNE outperforms t-SNE in preserving the global structure. The weight parameter λ of the global cost function determines the balance between local and global distances preservations. For the human brain atlas data set, we show the tradeoff of λ in representing global structure of data. Using g-SNE with the optimized λ may therefore yield biological insights into how data is organized on multiple scales. The MATLAB code is available at: https://github.com/gyrheart/gsne

## 1 Introduction

Dimensionality reduction techniques have been playing essential roles for analyzing modern high dimensional datasets. High dimensional data or objects, which are usually represented by high dimensional vectors or matrices of pairwise distances, can be embedded into lower dimensional spaces by preserving the data dissimilarities in the pairwise distances of embedded points. Low dimensional embeddings, for example in two or three dimensions, not only provide a way to visualize data organization but also reveal its hidden structure. The t-Distributed Stochastic Neighbor Embedding (t-SNE) is a powerful nonlinear embedding technique which had been widely applied in many areas of science, from visualizing feature representations in deep learning [1], to clustering bone marrow samples to distinguish between cancerous and healthy cells [2] and classifying neuron cells by gene expression profiles in biology [3]. As a neighbor embedding algorithm, t-SNE preserves the pairwise similarities of probable neighbors by minimizing the divergence of the similarity distributions between the neighboring data points and embedding points in the low dimensional space [4]. The similarity distributions describe the pairwise similarities of points, which peak at small distances but quickly decay to zero at large distances. The neighbor embedding property makes t-SNE effective for identifying local clusters in the data, but the side effect is that it fails to preserve the global inter-cluster structure – the embedding distances among clusters are meaningless and the global distribution of clusters is random [5]. The global structure of local clusters can provide significant insights into many biological systems. For example, ordering of cell clusters at different stages was found to represent a developmental trajectory [6] and to yield insights into cell lineages in the vertebrate brain [7]. For these tasks, it is essential to preserve inter-cluster organization of the data. Here we propose the global t-SNE (g-SNE) algorithm based on traditional t-SNE to preserve the global structure of clusters in data. We test the algorithm on synthetic dataset and two real datasets, and prove its ability to preserve both local and global structures in data.

## 2 Related work

After introducing t-SNE in 2008, Maaten et al. [4] further proposed improved version of t-SNE to speed up the algorithm using tree-based algorithm to approximate gradients [8]. The controllable t-SNE approximation was introduced to tradeoff speed and accuracy for interactive data exploration [9]. In biology, viSNE [2] was introduced to avoid crowding problems in low-dimensional embedding and to better visualize high-dimensional cells data in low-dimensional maps. These algorithms still used the local cost function which only captures the intra-class small distances. The g-SNE algorithm that we propose here is designed to improve how global structure is preserved. To achieve this, we introduce a global cost function component that is designed to capture inter-cluster large distances.

## 3 Global t-SNE (g-SNE)

In a data set containing *N* data points described by *D* dimensional vectors: {x_1_, x_2_, x_3_,…, x_*N*_; x_*i*_ ∈ ℝ^*D*^}. The t-SNE algorithm [4] describes the similarities of two points according to the following measure:

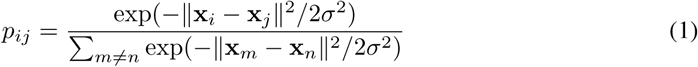

To avoid the crowding problem in the low-dimensional embedding, the heavy tailed Student t-distribution is applied within the embedding *d*-dimensional space where distances between points {y_1_, y_2_, y_3_,…, y_*N*_; y_*i*_ ∈ ℝ^*d*^} are defined as:

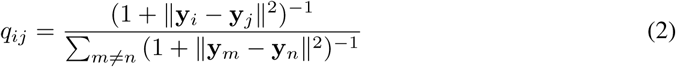

The Kullback-Leibler (KL) divergence between the joint probability distributions of pairwise data points *p_ij_* and embedding points *q_ij_* measures the distance discrepancies between the data and embedding points:

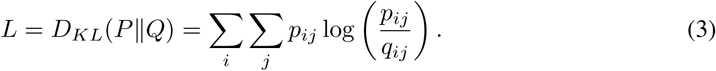

The t-SNE minimizes the KL divergence by gradient descent method. The gradient of the cost function *L* with respects to embedding coordinate y_*i*_ is (see Methods):

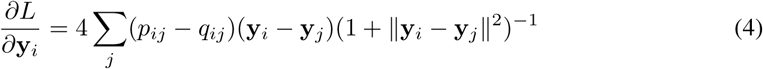

The probability distribution of distances *p_ij_* and *q_ij_* in Eqs. (1-2) are symmetric distributions that peak at 0 and decay exponentially or polynomially as the distances increase. As such, these distributions are sensitive to small pairwise distances among neighboring points but not to large distances of distant points. Minimizing the differences of probability distributions of data and embedding points can effectively capture and preserve local structure of data and generate well-separated clusters in embedding space. However, it fails to capture the large distances of inter-cluster points, so both the relative and absolute positions of the clusters are randomly embedded and meaningless [5]. To capture and preserve the global structure of the points and clusters, we propose g-SNE algorithm taking into account a new set of probability distributions for distance measures 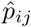 and 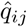 which are sensitive to large values but not to local ones:

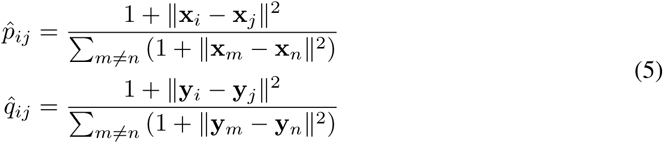

The global probability distributions 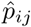 and 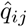 are also symmetric but peak at large values. Just like in Eq. (3), we can also define the global cost function 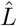 in g-SNE:

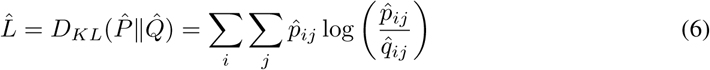

Minimizing the global cost 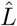 preserves the large distances in the low-dimensional embedding. To account for both the local and global structure of the data, we define a total cost function *L_total_* by combining the two cost functions using a weight parameter λ:

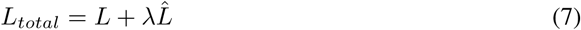

The gradient of the total cost function *L_total_* in g-SNE has a simple form:

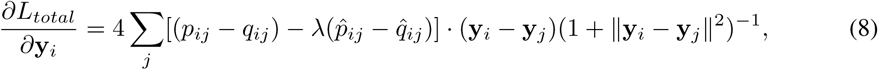

where the weight λ of the global cost function controls the balance between the local clustering and global distribution of the data. Large λ values lead to more robust global distribution of clusters, but less clear classifications. Small λ moves back to approximate the traditional t-SNE, and will be exactly the same when λ = 0. In next section, we will apply the g-SNE algorithm to one synthetic dataset and two real datasets to test its ability to preserve local and global structure of data, and compare that with t-SNE.

## 4 Experiment

Traditional t-SNE is powerful in generating local clustering, and it performs best in tasks where one only needs to define clusters, for example, to separate features in deep learning[1] but does not need pay attention to organization between clusters. To evaluate our new algorithm, we select the datasets where the global structures are significant and meaningful.

### 4.1 Snythetic data

We generate six groups of points in two-dimension plane, each group containing 50 clustered points; the six groups are hierarchically distributed in the plane (first column in Fig. 1). Two dimensional multidimensional scaling (MDS) [10] mapping of data recovers the data structure well (second column). The t-SNE generates six tightly clustered groups, but the distributions of the six clusters are random across three repeats and not consistent with the data structure (third column). After applying g-SNE to the data with λ = 1, the six well-separated clusters in the three repeats recover the global structure of the data, in terms of both the orientation and relative pairwise distances of clusters (forth column). This test on synthetic data shows the ability of g-SNE to preserve the global structure of data.

**Figure 1:**
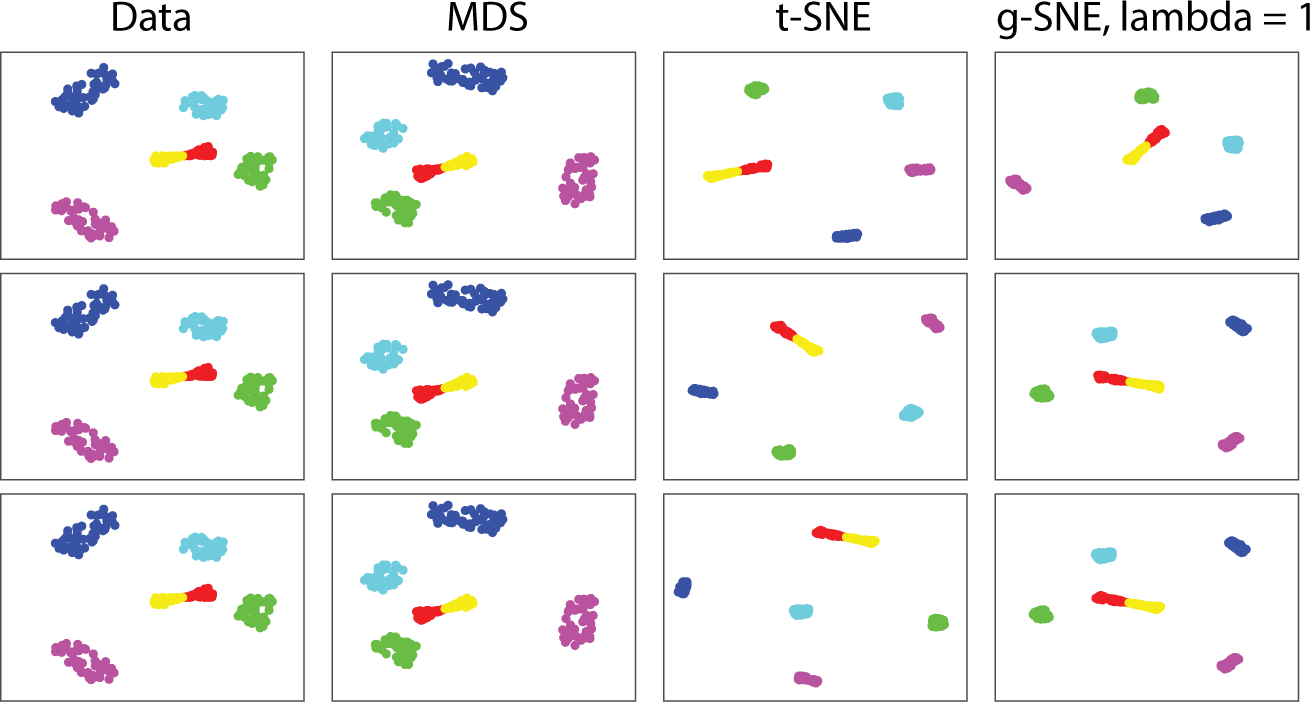
Clustering for synthetic data using MDS, t-SNE and g-SNE with three repeats. Three rows represent three repeats of mapping. First column: synthetic two-dimensional data containing six groups of points labeled by different colors, with 50 points in each group. Second column: MDS maps of data. Third column: t-SNE maps. Fourth column: g-SNE maps with λ = 1.

### 4.2 Iris flower data set

Next we apply the g-SNE algorithm to low dimensional real biological data set – the four-dimensional Iris flower data [11]. The Iris flower data set has been widely used in many statistical classification algorithms as a test example. It consists of 50 samples in each of the three *Iris* species: *setosa, virginica* and *versicolor*. Each sample is described by four features: the length and width of the sepals and petals measured in centimeters. We embed the four-dimensional *Iris* data set to two-dimensional space using MDS, t-SNE and g-SNE (Fig. 2). The 2D MDS mapping preserves the inter-cluster structure of the *Iris* data well across three repeats [12] (first colunm). The three species form three clusters, the *versicolor* cluster (green) and *virginica* cluster (blue) are close to each other and far from *setosa* (red), but *versicolor* is closer to *setosa* than *virginica*. The t-SNE mapping shows the three clusters, but the inter-cluster distances are not preserved: the *setosa* cluster is too far from the other two clusters compared with the MDS mapping, and the *versicolor* is farther from *setosa* than *virginica* in repeat 3 (second column). The g-SNE with λ = 1 generates very similar inter-cluster structure as the MDS in all three repeats, only with different rotations of the maps (third column). However, g-SNE provides better cluster separation than the MDS, which reduces the noise in the data. Therefore, the g-SNE combines both the advantages of the t-SNE in cluster separation and the ability of MDS to preserve the inter-cluster structure.

**Figure 2:**
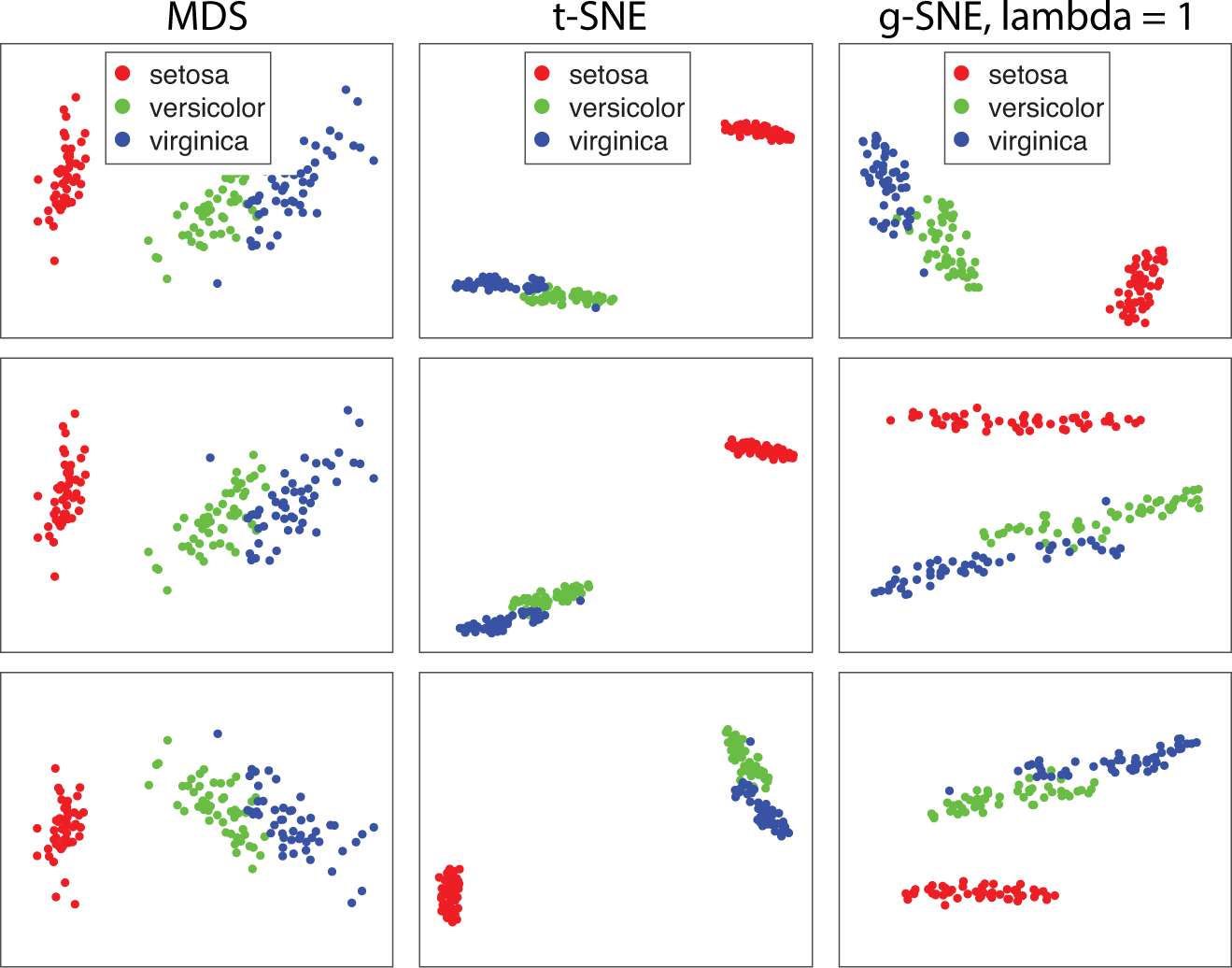
Clustering for *Iris* flower data set using MDS, t-SNE and g-SNE with three repeats. Three rows represent three repeats of mapping. First column: MDS maps of data. Second column: t-SNE maps. Third column: g-SNE maps with λ = 1.

### 4.3 Human brain atlas data

After successfully testing g-SNE on *Iris* data set, we now apply it to a much more complex and high-dimensional data – human brain transcriptome atlas [13]. This dataset contains microarray profiling of around 900 anatomical regions in human brain from two donors H0351.2001 andH0351.2002, and each sample region was profiled by 58692 probes representing 29191 genes. The data was already normalized using the methods in [14] and was available in [15]. Mahfouz et al. [3] applied BH-SNT (fast t-SNE) to human brain atlas data set to reduce the dimensionality of gene expression space and visualize the organization of region samples in the brain. Here, we first do the t-SNE mappings for the brain atlas data to reproduce the results in Mahfouz et al. [3]. We then apply g-SNE to the same data to generate new mappings and use recent topological methods [16] to quantitatively evaluate how well both methods preserve global structure of the data.

We apply both t-SNE and g-SNE to the gene expression profiles of brain region samples in donor H0351.2002 and plot the 2D maps of the samples with three repeats (first row in Fig.3A), the samples are colored by their anatomical acronyms from Allen Reference Atlas [15]. We identify the 15 acronyms used in [3] and label other acronyms as “Other” for comparison purposes. In total, we work with 16 clusters. To view the global structure more clearly, we calculate the mean positions of samples in each cluster (with same acronyms) as the centroid of the cluster, and plot the 16 centroids with the same colors (second row in Fig. 3A). The first repeat of t-SNE map resemble the result shown in [3]. However, the other two differ a lot in global distribution of clusters (Fig. 3A). The reason is that t-SNE performs weakly in preserving large distances in data. The g-SNE with λ = 1 shows different maps of the data, and the global structures of embedding clusters seem to be more consistent across three repeats than t-SNE maps (Fig. 3B).

**Figure 3:**
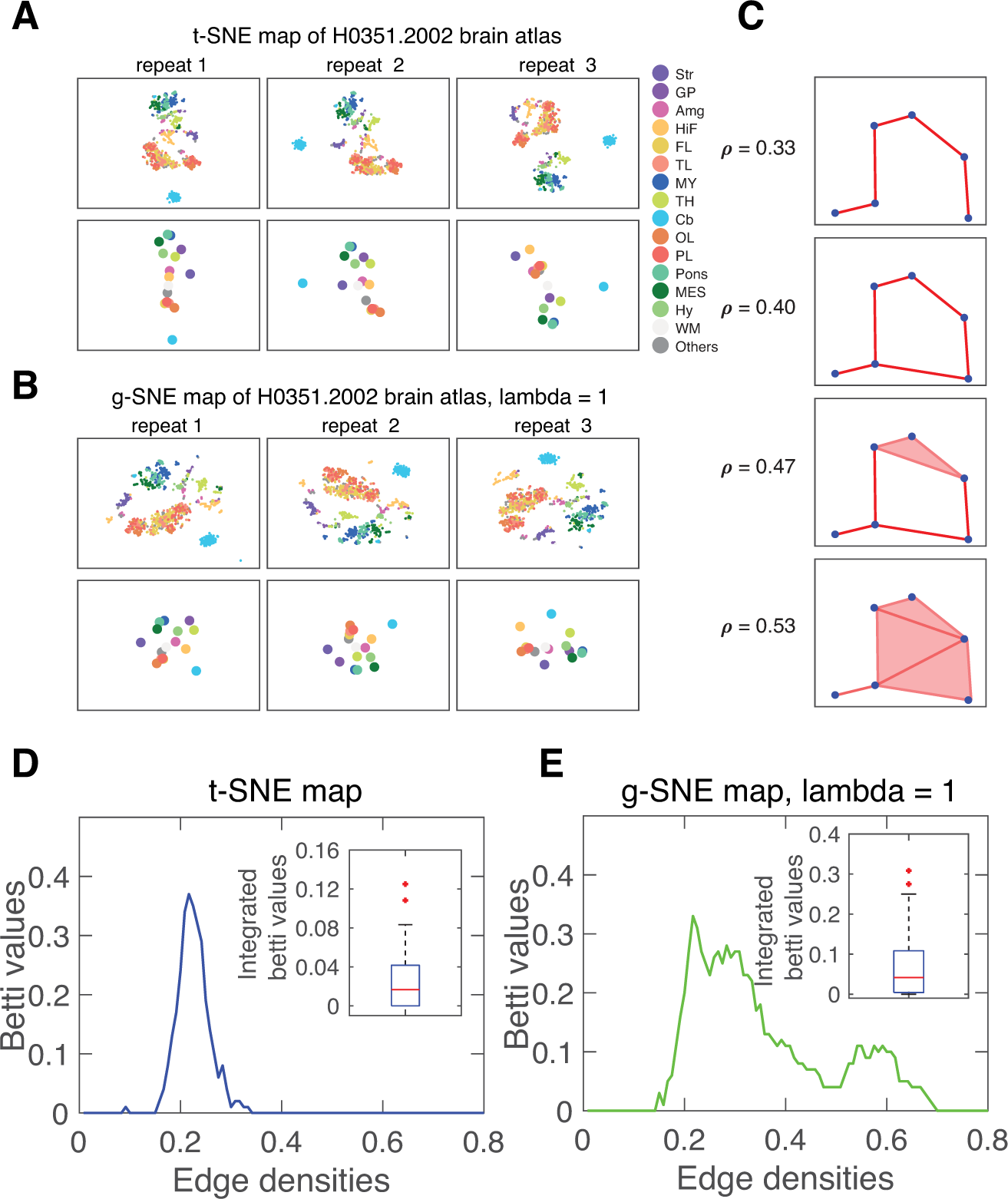
Two-dimensional embedding of gene expression profiles of donor H00351.2002 using t-SNE and g-SNE with three repeats. The brain region samples are labeled and colored by their anatomical acronyms from the Allen Reference Atlas: frontal lobe (FL), parietal lobe (PL), temporal lobe (TL), occipital lobe (OL), hippocampal formation (HiF), striatum (Str), Globus pallidus (Gp), amygdala (Amg), thalamus (TH), hypothalamus (Hy), mesencephalon (MES), Pons, myelencephalon (MY), cerebellum (Cb), white matter (WM) and Others. (A) Three repeats of t-SNE maps (first row) and the centroids positions of samples with the same labels (second row). (B) Three repeats of g-SNE maps with λ = 1 (first row) and centroids of the 16 anatomical groups in the maps (second row). (C) The schematic of network topology changing with the edge densities. New edges are added to the network as we decrease the connection threshold of points. The edge density *ρ* increases from 0.33 to 0.53, and the one-dimensional hole appears at *ρ* = 0.40, persists at *ρ* = 0.47 and vanishes at *ρ* = 0.53. Plotting the number of one-dimensional holes (first Betti values) against the edge densities yields the Betti curves in (D-E). (D) Average Betti curve of the 16 centroids in 100 repeats of t-SNE maps. Insets: box plot of integrated Betti values of the 100 repeats. (E) Average Betti curve of the 16 centroids in 100 repeats of g-SNE maps. Insets: box plot of integrated Betti values of the 100 repeats.

To quantify the global inter-cluster structure of the maps, we apply the topological technique from Giusti et al. [16] to characterize the distributions of cluster centroids in the 2D maps. According to the method, each pairwise distance matrix of a set of points can be quantified by a characteristic curve called Betti curve. The Betti curves are based on computing Betti values, which represent the number of holes of different dimensions with the set of connected points. The zeroth Betti value measures the number of connected components, whereas the first Betti value yields the number of one-dimensional “circular” holes. Points in the dataset are deemed “connected” if the distance between them is less than a certain threshold. Varying this threshold changes the number of connected data points and also affects the Betti value (Fig. 3C). The Betti curve describe how the Betti value changes as a fraction of connected points increases. Giusti et al. [16] reported that the integral of the Betti curve, termed the integrated Betti value, was sensitive to the presence of geometrical organization in the dataset, and in particular could distinguish geometrically generated data from random datasetss. For the tasks at hand we find that just working with the first integrated Betti value is sufficient to evaluate the data set structure.

To use the topological method, we generate the average Betti curves of the 16 cluster centroids from 100 t-SNE maps repeats, and make the box plots of the integrated Betti values of the 100 maps (Fig.3D). We make the same plots for 100 g-SNE maps in Fig.3E. As expected, the Betti curves of t-SNE maps and g-SNE maps have different shapes, and the integrated Betti values distribution of g-SNE maps is significantly different from t-SNE maps (*p* < 0.001 in two-sample Kolmogorov-Smirnov test). They obviously represent different global structures. To evaluate these two different results, we next explore the global structure of the gene expression data.

To obtain 16 representative points of clusters from the expression data, we cannot directly average the gene expression values of samples in the same clusters. One reason is that it may average out lots of important information, another reason is that it only produces one set of points and one Betti curve, which cannot be used to make statistical comparison with the results of 2D maps. To make full use of the data and generate a large number of inter-cluster representatives of Betti curves, we randomly select 16 anatomical samples from the 16 acronym groups, and then repeatedly sample different representatives from the 16 groups for 1000 times (Fig. 4A). As a control, we randomly take 16 samples from the whole brain but not based on acronym groups(Fig. 4B). After taking the samples, we can define pairwise distance matrix by the Euclidean distance of gene expression vectors, and then plot the Betti curves. We show the averaged Betti curves and integrated Betti value distributions of samples taken by acronym groups (Fig. 4A) and taken randomly across the whole brain (Fig. 4B). The difference of the two integrated Betti value distributions is significant (Fig. 4D, *p*<0.001 in two-sample Kolmogorov-Smirnov test). This shows that the data set has significant inter-cluster global structure. Integrating the Betti curves of data and 2D maps together shows that the Betti curves of g-SNE map better fits the data than t-SNE (Fig. 4C), and the integrated Betti value distributions of data is significantly different from t-SNE maps (*p* < 0.001) but not from g-SNE with λ = 1 (*p* = 0.25, Fig. 4D). Thus, g-SNE better preserves the global structure in data.

**Figure 4:**
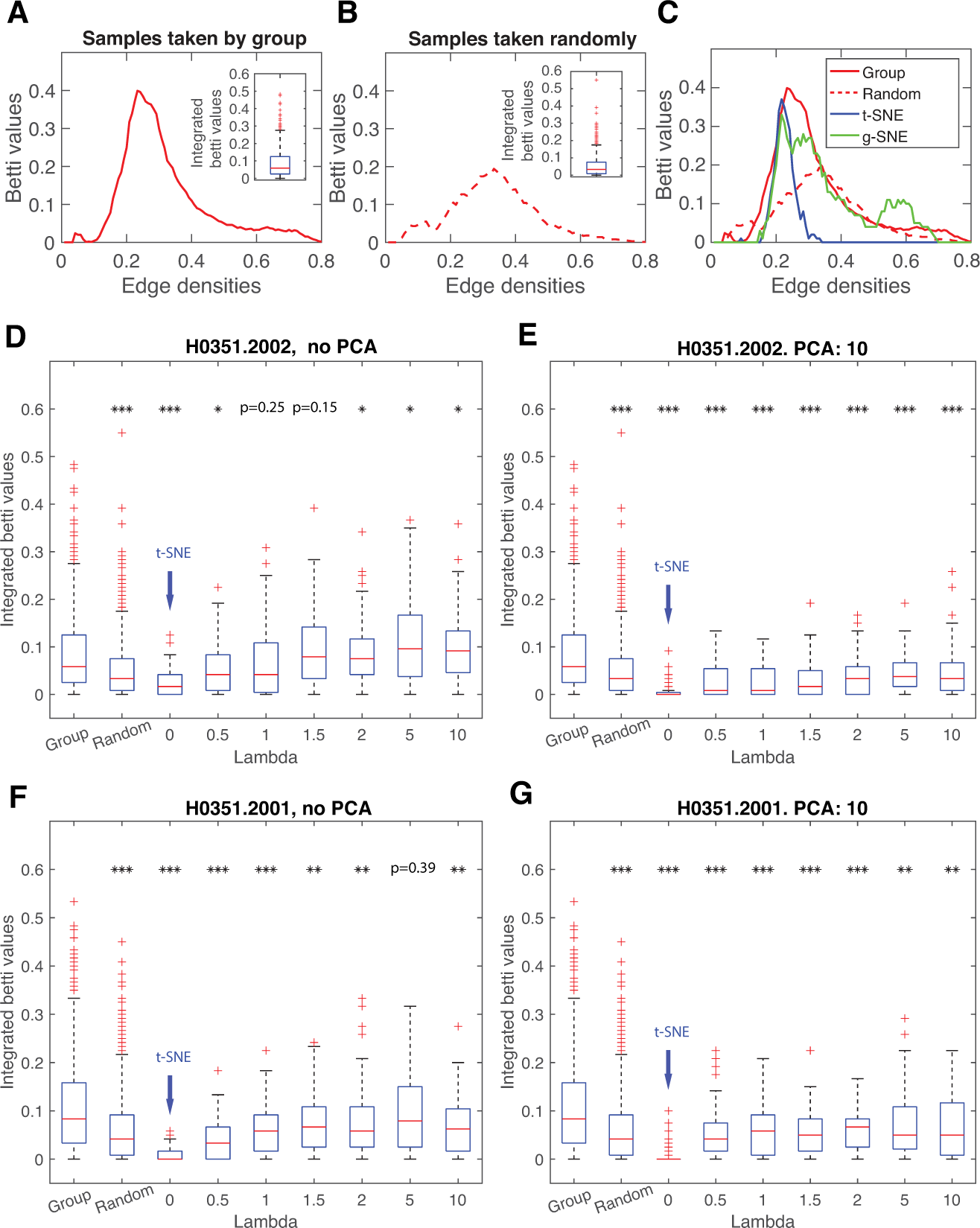
Betti curves of gene expression profiles of human brain atlas and evaluation of 2D embedding maps. (A) Average Betti curves of 1000 pairwise distance matrices, each matrix is generated from 16 anatomical region samples where each sample is randomly taken from one of the 16 acronym groups. (B) Average Betti curves of 1000 pairwise distance matrices, each matrix is generated from 16 samples randomly taken from all the anatomical samples in the brain. The pairwise distances of samples are defined as Euclidean distances of gene expression vectors. Insets showed the box plots of 1000 integrated Betti values of the Betti curves. (C) Integration of Betti curves of samples randomly taken based on acronym groups (red solid lines, “Group”), Betti curves of samples randomly taken from whole brain (red dashed line, “Random”), Betti curves of cluster centroids of t-SNE maps (blue lines) and Betti curves of cluster centroids of g-SNE maps with λ = 1 (green lines). (D) Box plots of integrated Betti values of data and g-SNE with different λ, λ = 0 is equivalent to t-SNE. No prior dimensionality reduction by PCA is done in g-SNE maps. (E) Box plots of integrated Betti values of data and g-SNE with different λ. Using 10 principal components of PCA for prior dimensionality reduction before doing g-SNE. Brain donor H0351.2002 is used in (A-E). (F-G) Box plots of integrated Betti values without (F) or with 10 principal components (G) of PCA as prior dimensionality reduction before g-SNE maps for brain donor H0351.2001. The stars in (D-G) represent the significance levels of two-sample Kolmogorov-Smirnov test on integrated Betti value distributions of the first column (“Group“) with the rest ones: *p* < 0.05(*), *p* < 0.01(**), *p* < 0.001(* * *).

In previous calculation, the parameter λ of g-SNE was set to 1. To further explore the properties of the algorithm, we screen other values of λ and plot the integrated Betti value distributions. We find that both λ = 1 (*p* = 0.25) and λ = 1.5 (*p* = 0.15) can fit the data, but either too small (λ < 0.5) or too large (λ > 2) cannot (Fig. 4D). The reason may be that too small λ act like t-SNE and cannot account for large distance distributions, while too large λ perform poorly in local clustering, which weakens the inter-cluster structure. So there is a tradeoff in λ in preserving global structure of data, and the optimal λ could provide best map for data. In Mahfouz et al. [3], they augured that higher order principal components helped to preserve the global structure, and that the t-SNE without prior dimensionality reduction by PCA did better in preserving global structure than that with PCA. We plot integrated Betti values of g-SNE maps with prior dimensionality reduction by selecting 10 principal components of PCA, and find that none of the λ could fit the data (Fig. 4E), consistent with the argument of [3].

Finally, we applied the same analysis to another brain donor H0351.2001. Here, both the integrated Betti value distribution of data and the optimal value λ = 5 (*p* = 0.39) were different from the previous person (Fig. 4D,F). The different optimal λ values of the two brain donors show that tradeoff between local and glocal structure can be case dependent. The g-SNE with prior PCA cannot fit the data (Fig.4G), which is the same as in the case of the previous person.

## 5 Discussion

We introduced the g-SNE algorithm by redefining the t-SNE cost function as the weighted sum of local cost function for local classification and global cost function for inter-cluster distribution. With this combined cost function, the g-SNE was able to preserve global structure of data as well as performing local clustering.

The weight parameter λ in g-SNE balances the local and global distances preservations. The t-SNE is a special case of g-SNE with λ = 0. When λ increases, the local distances constraint decreases while the global distances sensitivity increases. When λ becomes too large, the local clusters become hard to identify, thus the global structure also becomes less clear. Our experiment on brain atlas data illustrates the tradeoff in λ for preserving the global structure of data, and the optimal λ can be identified by comparing the integrated Betti values of cluster centroids in 2D g-SNE maps and anatomical samples taken from data.

When introducing the global joint probability distributions, we modified both the distribution functions of both data and embedding points as shown in Eq. (5). While this makes it possible to capture the global large pairwise distances, this measure may also become sensitive to the outlier points in data which may produce abnormally large distances and affect the embeddings. One way to solve the outlier-sensitivity problem is to change the power index *κ* of data distance metric in Eq. (5):

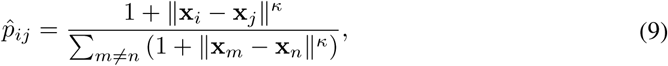

where *κ* ∈ [0,2]. Reducing the power index *κ* can reduce the sensitivity to outliers in data, but may also compromise the global structure preservation. Therefore, the power index may need to be tuned and optimized based on the quality of the data.

When applying g-SNE on another data set (H0351.2001), the optimal λ changes, indicating that optimal λ is case dependent. How the optimal λ is related to the intrinsic structure of data would be an interesting problem to study in the future. In particular, it may provide new approaches to identifying the statistical differences across different data sets with similar data structure, for example, finding the human transcriptome differences across tissues and individuals [17]. Identifying global structure in the data has great applications in biology, for example, it helps to infer cell lineages from gene expressions [7]. It is our hope that the g-SNE algorithm would help in these endeavors.

